# The elephant retina examined across a range of ages

**DOI:** 10.1101/2021.01.20.427452

**Authors:** James Freed, Saskia Joyce, Rachel Whalley, Edward Purse, Thomas L Ingram, Lisa Yon, Lisa Chakrabarti

**Affiliations:** School of Veterinary Medicine and Science, University of Nottingham, Sutton Bonington, LE12 5RD, UK; MRC Versus Arthritis Centre for Musculoskeletal Ageing Research, UK

## Abstract

The effect of aging in the human retina has been well documented, as have the signs of age-related retinal disease. Comparative studies in animals allow us to further investigate how the retina ages in different species. The African bush elephant (*Loxodonta africana*) has a retina comparable to other mammalian species, but with some reported distinctive differences in retinal ganglion cell (RGC) distribution and type. This is a first survey of the retina of *Loxodonta africana* from individuals aged 2 months to 32 years old. Gross examination, H&E staining and cell counts were used to compare calves (0-5 years), juveniles (6-10 years), sub-adults (11-20 years) and adults (>20 years). Dorsal-ventral eye diameter was shown to be significantly greater with age, whilst no significant changes in photoreceptor number were found. Changes in retinal thickness differed from past findings in elephants and humans, with thickness decreasing, then increasing in adults. Various morphological differences were evident in the samples including the presence of nuclei in the outer plexiform layer (OPL), degeneration of the inner plexiform layer (IPL), interruptions in nuclei columns of the outer nuclear layer (ONL), and larger unidentified cells within the inner nuclear layer (INL). These are initial observations provide some baseline information for a species where this range of samples has not been described previously.

## Introduction

The vertebrate neuroretina is composed of five layers: three layers of nerve cell bodies and two layers of synapses. Photoreceptor cell bodies compose the outer nuclear layer (ONL) and bipolar, horizontal, and amacrine cell bodies are situated in the inner nuclear layer (INL). Ganglion cell bodies and some amacrine cells make up the ganglion cell layer (GCL). The inner plexiform layer (IPL) and outer plexiform layer (OPL) are the synaptic clefts delineating the nuclear layers (Bonnel et al., 2003).

Aging can be defined as a gradual deterioration causing loss of physiological function which in turn increases the likelihood of disease (L?pez-Ot?n et al., 2013). Somatic mutations accumulate with age, resulting in altered proteins that reduce effective cellular function (Szilard, 1959; Gonzalez-Freire et al., 2015; Milholland et al., 2017). Cellular senescence is a mechanism that responds to ageing effects by arresting the cell cycle. Accumulation of senescent cells in aged tissues can be considered to contribute to dysfunction (Campisi, 2013). Cell senescence causes activation of TOR (target of rapamycin), leading to hyperfunction of cellular processes, then reduced protective mechanisms. (Blagosklonny, 2006, 2012).

The human retina changes with age, with decreased neuronal and retinal pigment epithelial (RPE) cells, increased lipofuscin within RPE cells, and increased thickness and drusen accumulation of the Bruch’s membrane (Newsome et al., 1987; Ramrattan et al., 1994). Cells within the GCL, and rod photoreceptor cells, decreased the most in the aged retina (Gao and Hollyfield, 1992). In the older retina, the width of the layers appeared unchanged, but cone and RPE cells demonstrated reduced density (Wang et al., 2020). A maintained retinal thickness through increased retinal area, despite thinning of layers, was seen in aged mice (Samuel et al., 2011). A failure of the photoreceptor to synapse with the bipolar and horizontal cells has been demonstrated in ageing human and mouse retina (Eliasieh et al., 2007; Samuel et al., 2011). Numbers and densities of rod and cone photoreceptors do not change in aged mice as they do in humans, but electroretinography findings are reduced, with reduced synapses the potential cause (Samuel et al., 2011; Sugita et al., 2020). Compensation has been described in aged mice and humans, where rod bipolar cells extend their dendrites into the ONL to synapse with rod photoreceptors (Liets et al., 2006; Eliasieh et al., 2007). Studies in birds and reptiles have shown lipofuscin accumulation in the RPE similar to humans, and deterioration of this layer, and photoreceptor layers of the retina, with similar changes reported in horses (Ehrenhofer et al., 2002; El-Sayyad et al., 2014).

Elephant retinal layers follow the general vertebrate pattern, but there is no fovea, and rod photoreceptors predominate (Kuhrt et al., 2017). Elephants, like many mammals, show a visual streak on the retina: a horizontal band of increased photoreceptor density. (Mullins and Skeie, 2010). However, the increased density of ganglion cells in the upper temporal retina is only seen in elephants and has been suggested to provide binocular vision when using the trunk (Stone and Halasz, 1989). The nasal area centralis is a third region of increased ganglion density which allows greater visualisation of the posterior visual field (Pettigrew et al., 2010). The elephant retina is paurangiotic (avascular), similar to that of the horse, while their soma size variability compares with rabbits and rats (Eorge L Indsay and Ohnson, 1901; Stone and Halasz, 1989; Ehrenhofer et al., 2002). Elephants display giant soma of retinal ganglion cells (RGC) which are comparable to those of the dolphin and could be a means of compensating for the lower number of RGC when compared to other mesopic mammals (Dawson et al., 1982; Kuhrt et al., 2017). Their rod bipolar cells appear to be bistratified, whereas in other mammals they are monostratified. Elephants are active both day and night, which may explain why their rod bipolar cells are bistratified, similar to Microbats, as this can help with nocturnal contrast perception in low light (Pettigrew et al., 2010; Müller et al., 2013).

Few studies have been conducted on the elephant retina, and even fewer have had access to such a range of ages as those assessed for this study. African bush elephants live for a maximum of 70-75 years, whilst average lifespan for humans is 72 years (Lee et al., 2012; Life expectancy at birth, 2015). Looking at age-related changes in elephant retina could therefore be interesting to compare with retinal aging in humans.

## Materials and Methods

### Collection of elephant retinal tissue

Collection of elephant retinal tissue was approved by the local ethics committee of the School of Veterinary Medicine and Science at the University of Nottingham. The samples were obtained from management organised culled operations at Save Valley Conservancy (SVC), Zimbabwe between 2009 and 2011. The elephants were not culled specifically for this study and permission was acquired from all pertinent authorities. The Zimbabwe Parks and Wildlife Management Authority (PWMA) gave permits to SVC to cull the elephants and the SVC gave permission for the samples to be used by the School of Veterinary Medicine and Science. The eyes were collected from each of the 24 elephants (see Table 1). One eye from each pair was immersed in liquid nitrogen, shipped to the United Kingdom and stored at −80°C. The other eyes were preserved and immediately shipped to the UK in 10% neutral buffered formalin. These eyes were then immersed in 4% paraformaldehyde (PFA) in phosphate-buffered saline (PBS), which was then later reduced to 0.4% PFA.

**Table 1:**
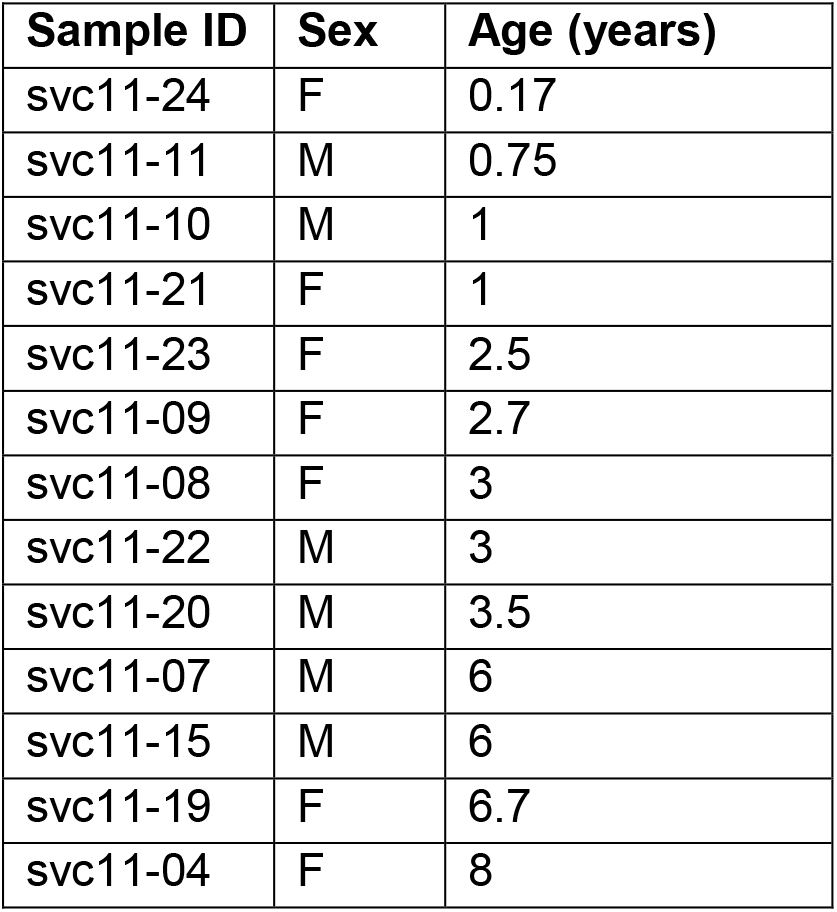
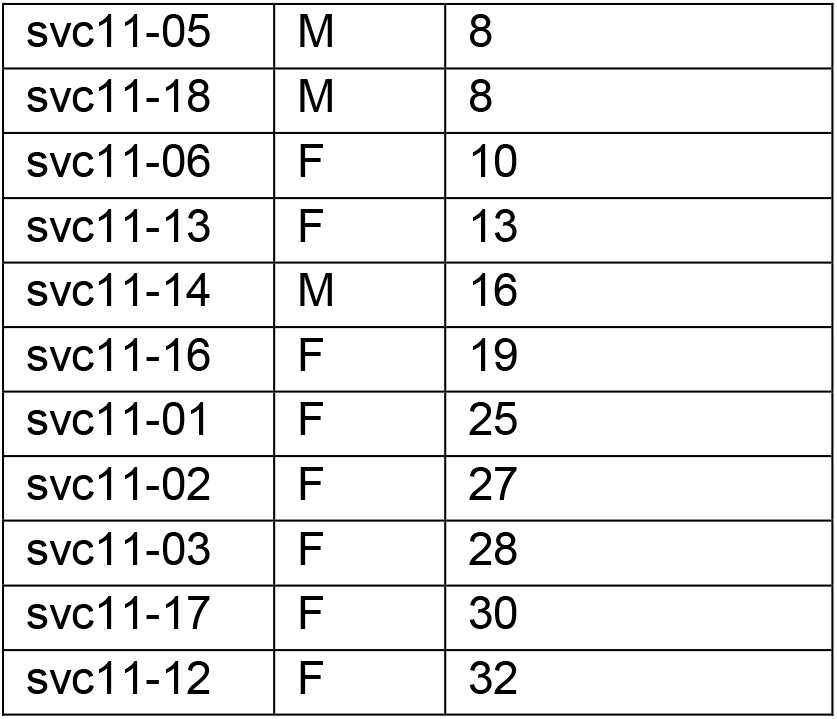
Elephant identification number, sex and age. Ages range between 2 months and 32 years. Males and females (n=9 and n=15 respectively) were not age-matched, and therefore differences in retinal structure between sexes were not investigated.

| Sample ID | Sex | Age (years) |
| --- | --- | --- |
| svc11-24 | F | 0.17 |
| svc11-11 | M | 0.75 |
| svc11-10 | M | 1 |
| svc11-21 | F | 1 |
| svc11-23 | F | 2.5 |
| svc11-09 | F | 2.7 |
| svc11-08 | F | 3 |
| svc11-22 | M | 3 |
| svc11-20 | M | 3.5 |
| svc11-07 | M | 6 |
| svc11-15 | M | 6 |
| svc11-19 | F | 6.7 |
| svc11-04 | F | 8 |
| svc11-05 | M | 8 |
| svc11-18 | M | 8 |
| svc11-06 | F | 10 |
| svc11-13 | F | 13 |
| svc11-14 | M | 16 |
| svc11-16 | F | 19 |
| svc11-01 | F | 25 |
| svc11-02 | F | 27 |
| svc11-03 | F | 28 |
| svc11-17 | F | 30 |
| svc11-12 | F | 32 |

### Dissection

The eyes were trimmed to remove surrounding fat and optic nerve tissue. The front of the eye was dissected at the coronal plane of the globe. The vitreous humour, aqueous humour and lens were then removed. The eye was dissected from dorsal to ventral, through the optic nerve, resulting in a section containing both tapetal and non-tapetal fundus. A second section was made from dorsal to ventral, creating a cross section of the eye. The sections were then placed in cassettes containing 70% ethanol until ready for processing.

### Processing and embedding

Sections were embedded in paraffin wax as described in Werner et al., (2000). The tissue was dehydrated in 70%, 95% and 100% ethanol, followed by the use of histoclear to wash the tissue. The cassettes were then placed in paraffin with the optic nerve edge facing downwards and left at 4°C to solidify.

### Tissue sections

For H&E staining, cross-sections of the retina were sectioned at 8μm using a fully automated rotary microtome Leica RM2255 (Leica Microsystems, UK) and placed onto Superfrost™microscope slides (Thermo Scientific, UK).

### H&E stain

Five cross-sections of each eye were stained as described in Fischer et al., (2008) with the following changes. The slides were deparaffinised in Histo-Clear II (National Diagnostics, UK) twice for 5 minutes, then rehydrated in 100%, 95% and 70% ethanol for 2 minutes each. Slides were washed in running water for two minutes. Haematoxylin (Sigma-Aldrich, UK) was added to the slides for 5 mins. Between the addition of haematoxylin and eosin, 1% acidified IMS and 50% ammoniated water were used with running water between steps. Eosin (pH 5.0) (Sigma-Aldrich, UK) was used for 5 minutes with running water, followed by dehydration using intervals of 70%, 95% and 100% ethanol. Slides were washed in Histo-Clear II twice for 5 minutes each, then submerged in Xylene (Fischer Scientific, UK) for 5 minutes.

## Calculations

Axial eye diameter was measured from the anterior pole of the eye to the posterior pole (mm). Dorsal-ventral diameter was measured from the central eye (mm). Photoreceptor counts were determined as shown in figure 1, and an average for each retina calculated. Retinal layer thicknesses were measured in micrometres by light microscopy at 20μm. The tapetum of each sample was divided into one of 4 categories depending on colour: All blue (1), predominately blue with some white (2), predominately white with some blue (3), and all white (4).

**Figure 1:**
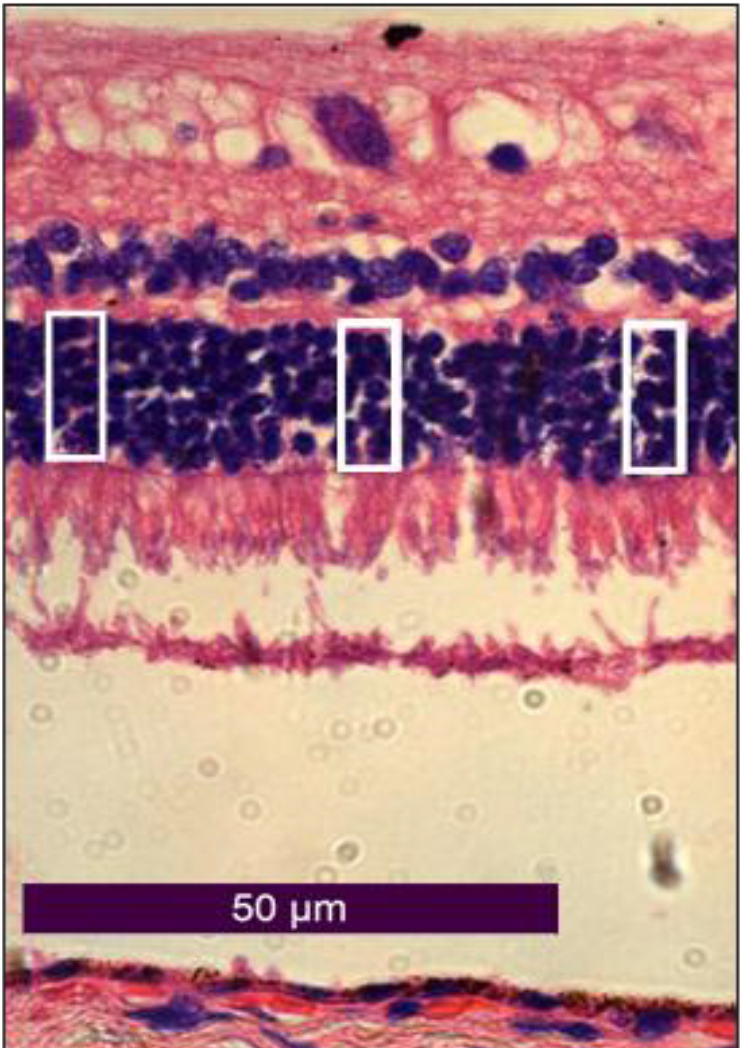
White boxes show columns counted with 5 photoreceptor nuclei in ONL. H&E stain of svc11-06. Section 8μm thick. 400× magnification.

**Figure 2:**
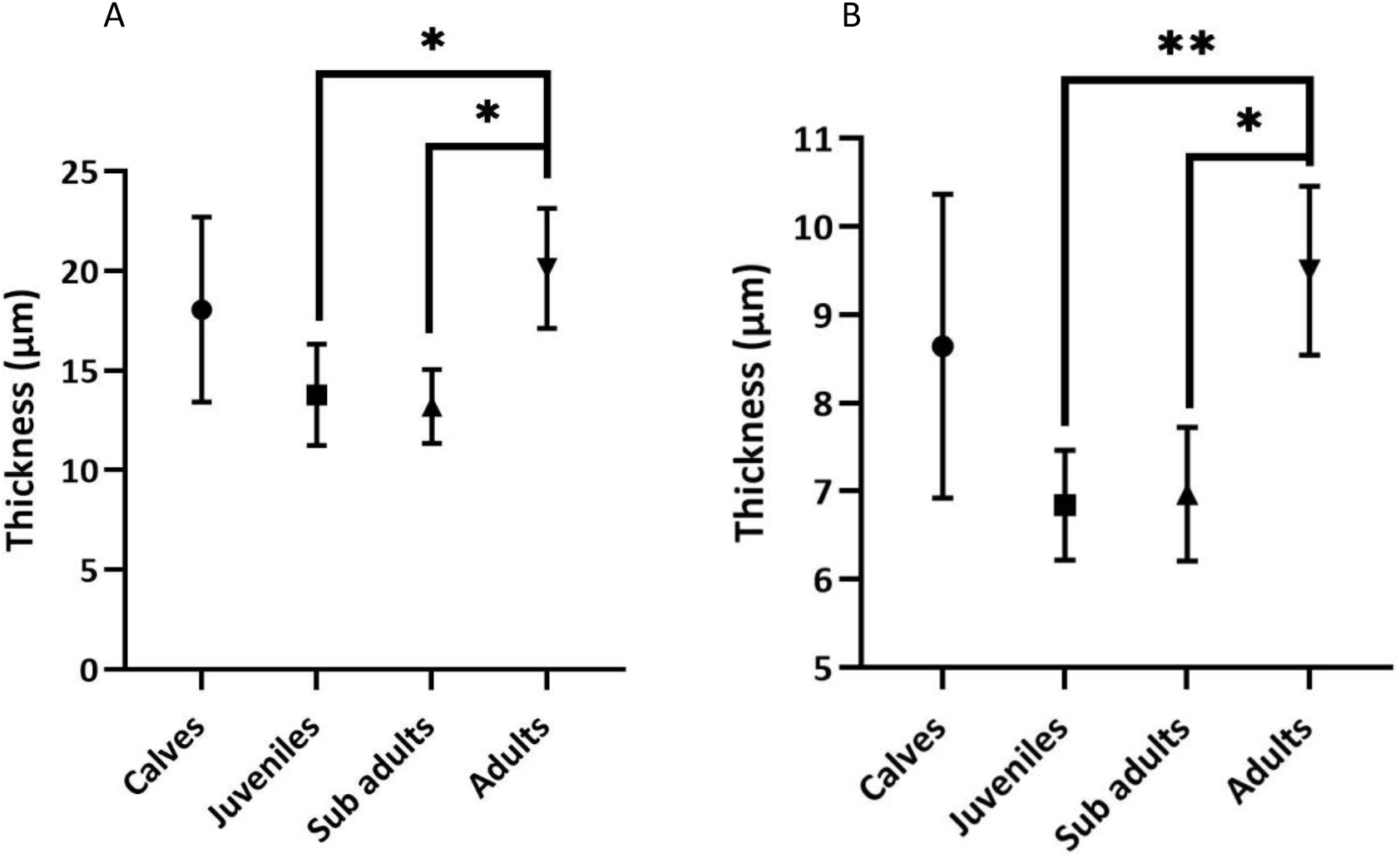
Graphs showing means and range of the ONL (A) and INL (B) of the retina. Groups were calves (0-5), juveniles (6-10), sub-adults (11-20) and adults (>20). Significance bars are shown.

## Results

Data were grouped by age according to ranges defined in Stoeger et al., (2014).

## Discussion

This study arose from a unique opportunity to obtain and examine a number of non-captive elephant eyes across an age range spanning several decades. The elephants were from a single group which assured a homogeneous environmental background. We aimed to see whether there were clear age-related changes in the elephant retinas that were collected.

We show that the dorsal-ventral diameter of adult African elephant eyes was significantly larger than that of the younger groups. Between calves and adults, the diameter increased by a factor of 1.19. The axial diameter between calf and adult also significantly increased, by a factor of 1.13. These findings are suggestive of growth through aging (Kuhrt et al., 2017), and correlates with previous work which determined that eye mass increases with body mass (Howland et al., 2004). All age groups had a dorsal-ventral diameter greater than the eye’s axial diameter (Table 2). Both African and Asian infant elephant eyes have been demonstrated to be more spherical than the adult, as seen here (Murphy et al., 1992; Bapodra et al., 2010; Pettigrew et al., 2010). When the diameters are compared to those of Asian elephants, the African elephants’ eyes have been reported to be larger (Bapodra et al., 2010). We found larger values for both axial and dorsal-ventral eye diameter in African elephants when compared to their Asian counterparts (Table 2).

**Table 2:** Dorsal-ventral eye and axial diameter increases significantly from calf to adult. The mean axial and dorsal-ventral diameters for calves (0-5), juveniles (6-10), sub-adults (11-20) and adults (>20). Elephant eye samples svc11-04, −07, −08 and −10 were excluded as they had already been processed. Using a one-way ANOVA, we found that there is a significant difference in dorsal ventral diameter between groups (p=0.031). Using an unpaired t test, we found a significant difference in the axial diameter when calves were compared to adults (p=0.04).

| Measurement | Calf | Juvenile | Sub-adult | Adult |
| --- | --- | --- | --- | --- |
| Axial diameter (mm) | 32.29± 3.45 | 34.60 ± 8.99 | 35.33 ± 1.53 | 36.60 ± 2.61 |
| Dorsal-ventral diameter (mm) | 36.14± 3.85 | 38.40 ± 1.52 | 42.33 ± 5.51 | 43.00 ± 4.47 |

On gross examination of the eye, most tapeta observed were wholly or mostly blue (Figure 3). This concurs with findings from an infant African elephant, and is a trait shared with the cow, juvenile cat, dog, sheep and goat (Stone and Halasz, 1989; Ollivier et al., 2004). However, it has been shown that this is silver to white in the adult elephant (Pettigrew et al., 2010). There has been a suggestion that the colour of the tapetum changed with age in Asian elephants (Murphy et al., 1992), but the few elephants that did display a more white tapetum in our sample set were of various ages, with no clearly discernible pattern. In fact, in our study, the oldest elephant’s eye had a blue tapetum, and the one-year-old had a predominately white one. This difference has previously been attributed to the strength and location of the light source used when assessing the colour of the tapeta (Pettigrew et al., 2010).

**Figure 3:**
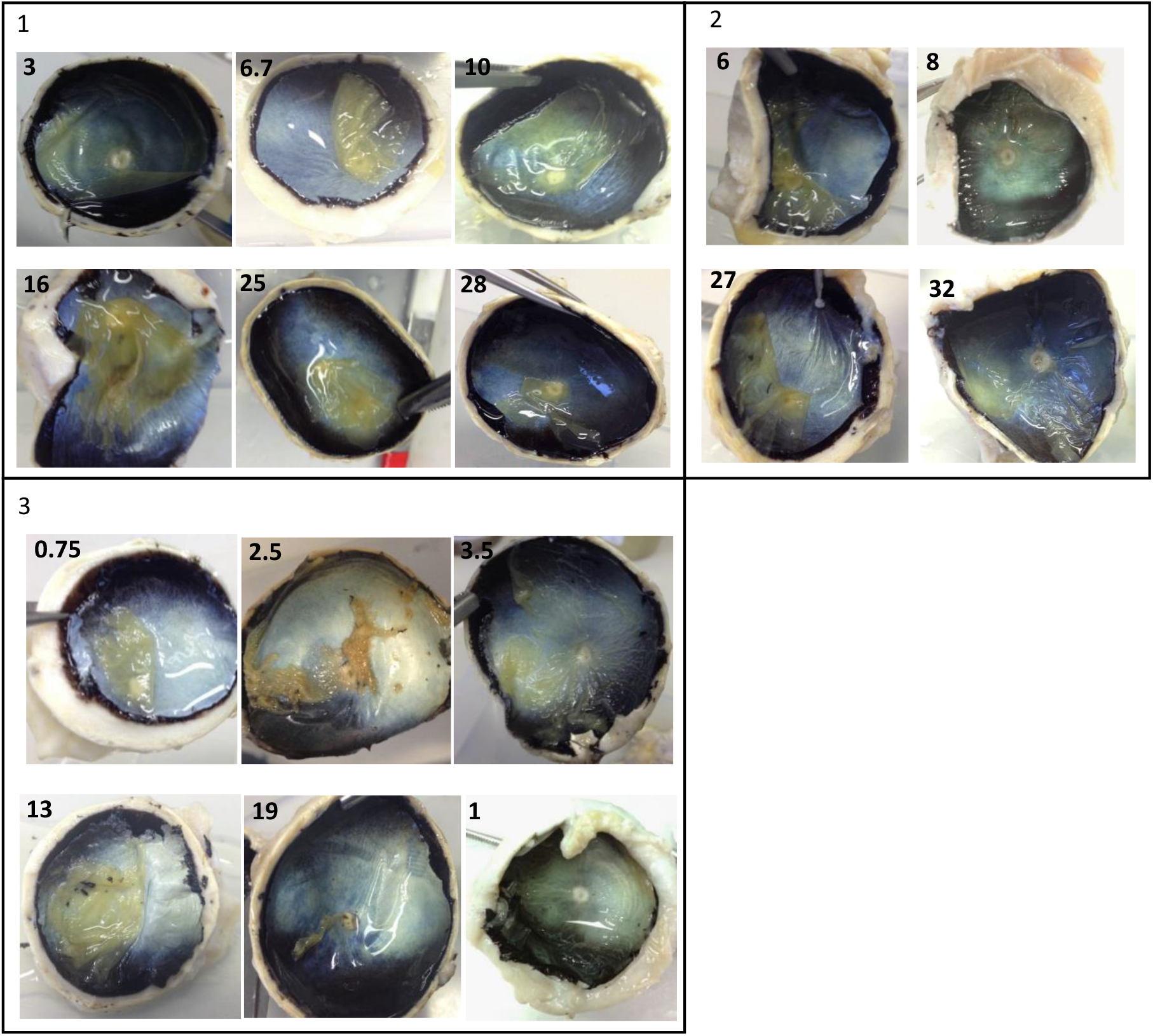
The tapetum colour showed no apparent age-related pattern. The posterior of the eye. Panel 1: Blue. Svc11-22, svc11-19, svc11-06, svc11-14, svc11-01, svc11-03 (left to right). 2: Predominately blue. Svc11-15, svc11-04, svc11-02, svc11-12 (left to right). 3: Predominately white. Svc11-11, svc11-23, svc11-20, svc11-13, svc11-19, svc11-10 (left to right). Age in years shown in top left of image. There was no image for svc11-07 and −18. Svc11-24 not included due to poor visualisation of the tapetum.

In the histological assessment, to enable comparison across the age groups, the area lateral to the optic disc was examined to ensure similarity in the layer composition. In cases where limited quantities of retina were captured in the slide, we used what was available. Unfortunately several of the retinas became detached during the process of collection and transfer to our laboratory, however, this is unsurprising, given what was involved in collecting these samples in the field in Zimbabwe. Indeed, the logistical challenges which had to be addressed to gather these samples at all is reflected in the paucity of currently published work on this subject.

The average photoreceptor count showed the ONL had between 4-5 nuclei per single layer cross section of retina (table 3). By contrast, in other studies in the African elephant, a functional column of retina has been shown to contain 11 photoreceptors in total, over double our count (Kuhrt et al., 2017). This may be down to difference in sample thickness, level of preservation, counts taken from different parts of the retina or a wider definition of a functional column. There was no significant difference in the number of nuclei between the younger and older elephants. This contrasts with a past study which reported that the photoreceptor density was higher in newborns than adults (Kuhrt et al., 2017). That same study found elephant retinas to be relatively mature at birth, having all layers and most cell types found in adults, which may be why our counts remained similar across the ages examined. In humans the fastest level of photoreceptor loss occurs around midlife, between the 2nd and 4th decade (Gao and Hollyfield, 1992; Curcio et al., 1993). The lack of degeneration in the elephant by this point may indicate that elephant eyes have mechanisms to maintain health, or deteriorate at a slower rate compared to humans. Alternatively, it could be that elephants with poor vision do not survive well in the wild, and therefore were not available for sampling. Total counts could have been calculated if specimens were all equally well preserved, in order to calculate photoreceptor density more accurately, and this is certainly an area for future study.

**Table 3:** Photoreceptor counts are not significantly different between the age groups. Photoreceptor count from the ONL in each of the 5 H&E slides for calves (0-5), juveniles (6-10), sub-adults (11-20) and adults (>20. All eyes containing retina were included, though counts could not be taken from slides that were overstained. From a one-way ANOVA test, we suggest that there is no significant difference in the number of photoreceptor nuclei between the *L. africana* retinas studied (p= 0.25).

Photoreceptor counts are not significantly different between the age groups
|  | Calf | Juvenile | Sub-adult | Adult |
| --- | --- | --- | --- | --- |
| Photoreceptor count | 4.35 | 5.05 | 4.6 | 4.4 |

The thickness of the photoreceptor dendrites and OPL were not significantly different between the age groups of these elephants. However, the difference in the ONL and INL thickness was significant. The ONL and INL thickness from our results (Table 4) differed from past counts, with both calves and adults having an ONL approximately a third thinner, and an INL roughly half as thick as compared to results reported from past research (Kuhrt et al., 2017). The numbers of photoreceptor nuclei we counted were also different from the Kuhrt et. al., study, which may be due to variations in the region of the retina sampled. To confirm that the variation in retinal layers and photoreceptors is a genuine finding, in future studies samples should be taken from the same area of the retina. In previous work, INL thickness was reported to decrease from newborn to adult in both African and Asian elephants. However while, ONL thickness decreased from newborn to adult in Asian elephants, in the African elephants ONL thickness was greater in adults, though the reason for this difference is unknown (Kuhrt et al., 2017). Our results differed from those previously reported, with the thickness of the ONL and INL decreasing from juvenile to sub-adult, but then increasing again within the adult age group. Density of cells and layer thickness of the retina in adults may decrease due to eyeball growth causing stretching and horizontal expansion of the retina, which occurs in many mammals (Kuhrt et al., 2012). This would explain the reduced thickness with age initially, although if this were the case, then we would expect that the adult thicknesses should be smaller still, which was not supported by our findings. Common causes of retinal thickening in humans are macular oedema and retinal fibrosis (Nussenblatt et al., 1987; Friedlander, 2007). Elephants lack a macula, instead having a visual streak (Stone and Halasz, 1989), and it is currently unknown if similar pathology can result in retinal thickening via the same mechanisms seen in humans. This would be a possible reason for thickening of the retina, but in our observations only the nuclear layers were thicker. The nuclei of human fibroblast cells have been shown to swell with senescence, which is caused by MAP kinase (Kobayashi et al., 2008), and MAPK inhibitors have been shown to be a potential therapeutic for AMD (Kyosseva, 2016). The adult elephants may have higher numbers of senescent cells that have swollen, leading to thickening of the ONL and INL. This was not seen in our study, but the oldest elephant sampled in this study was 32 years of age, which is less than middle-aged for an African elephant (Lee et al., 2012), so perhaps older elephants would have more apparent age-related changes.

**Table 4:**
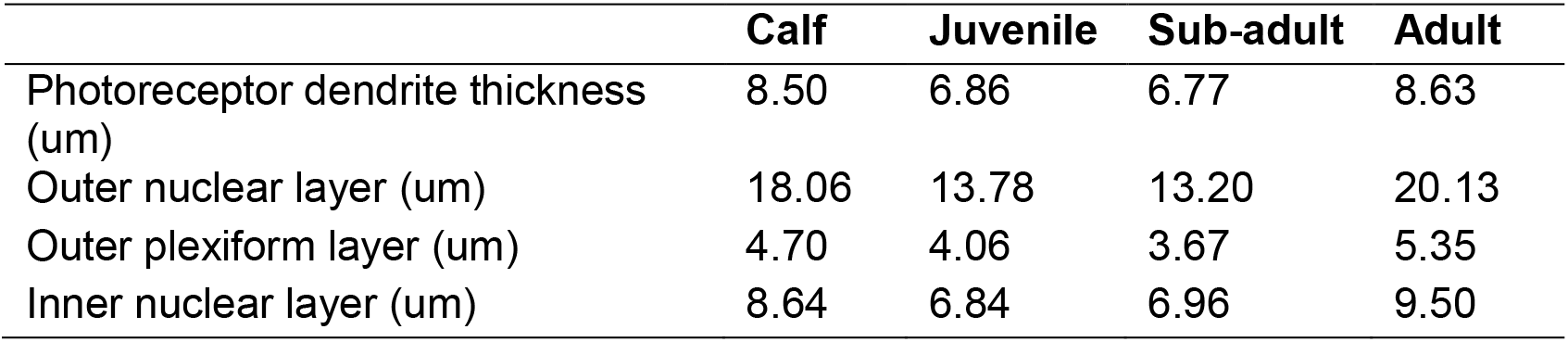
The thickness of the nuclear layers are significantly different between juveniles and adults. The mean thickness of the photoreceptor dendrite layer, ONL, OPL, and INL for calves (0-5), juveniles (6-10), sub-adults (11-20) and adults (>20). Counts were taken from all eyes containing retina, (Table 6). With an unpaired t-test, we found a significant difference between the ONL thickness of adults and juveniles (p=0.011), and between adults and sub-adults (p=0.018). With an unpaired t test, we found significant difference for the thickness of the INL between the adults and juveniles (p=0.0015), and also between adults and sub-adults (p=0.0133).

Many samples displayed cell bodies with nuclei in the OPL (figure 4), which is usually comprised of synapses alone. These might be the giant ganglion cells that are known to be present in elephant retina (Kuhrt et al., 2017). However we observe the cell bodies to be sited deeper into the layers, and their size did not match that of the other ganglion cells in the sample (figure 7). Photoreceptors can retract into the OPL in dyslamination in disease, and this causes depopulation of the ONL (Li et al., 2018). If disease were present, this may then also account for the missing nuclei within the ONL, also observed in the 28-year-old eye (figure 6). These large gaps were also seen in the oldest eye in this study, but no nuclei were observed in that ONL. Although the interruptions in the ONL may be due to age-related degeneration in the older eyes, it was observed in younger eyes as well. In the 2-month-old eye, it may have been the result of photoreceptor apoptosis as the retina developed, which has been shown to occur up to 72 days postnatally in the rat (Vecino et al., 2004). For the other age groups, it is known that dendrites of the photoreceptors spread into the ONL, suggesting that physical ingression between nuclei columns could cause them to separate from each other, as dendrites require space (Eliasieh et al., 2007). Deterioration of the IPL was observed (figure 5), which may be due to degeneration, or to physical damage which occurred when the samples were handled and prepared. However, trans-synaptic degeneration has been shown to occur between cell bodies in the ONL and INL in mice and humans with aging (Eliasieh et al., 2007; Samuel et al., 2011), so similar changes may occur between the GCL and INL. Patients with neovascular diseases such as AMD also have thinner IPL (Lee et al., 2020). The variable ages of the three samples showing IPL loss suggests it is unlikely to be an age-related change we see here.

**Figure 4:**
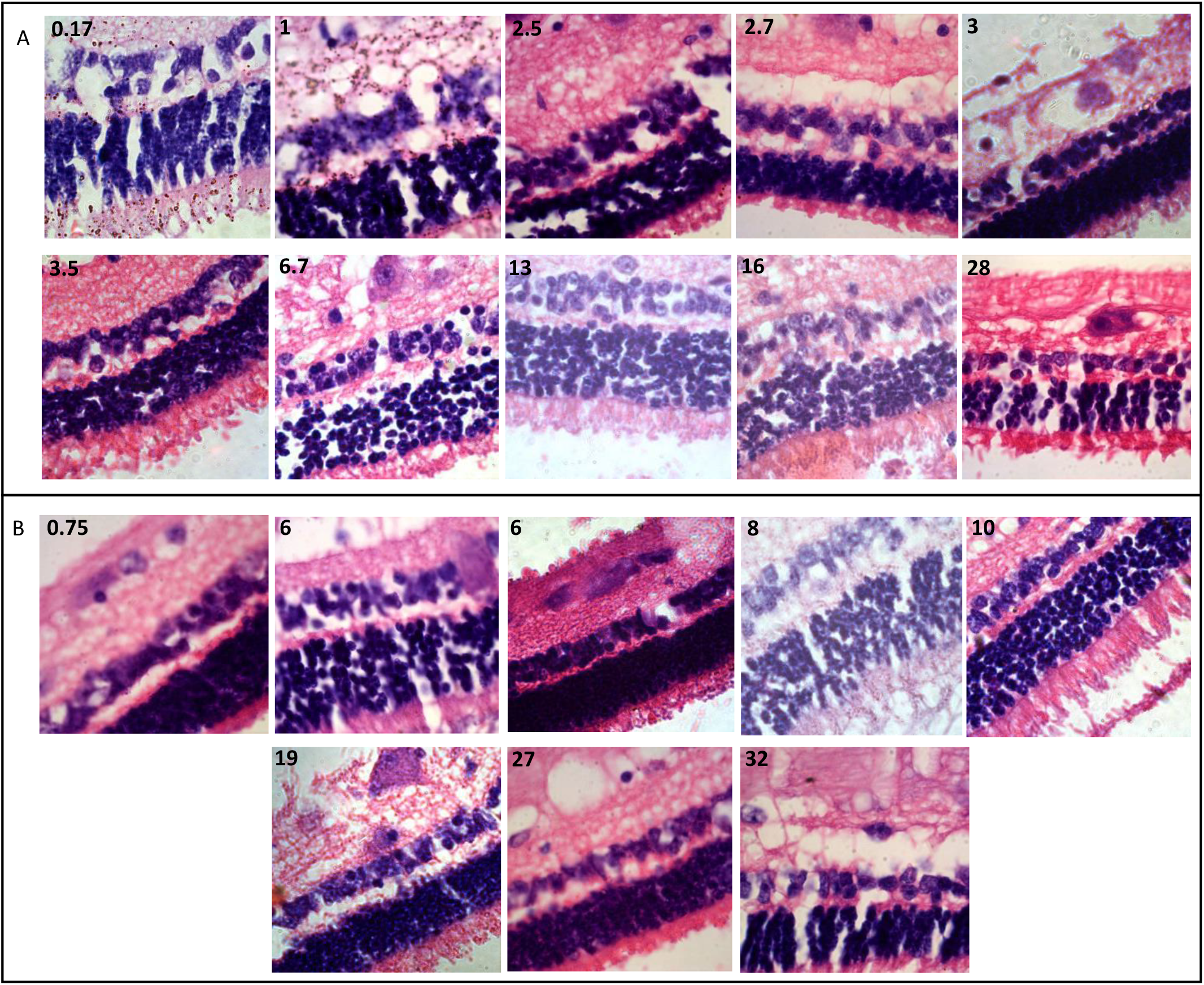
Nuclei were found in the outer plexiform layer of several elephant retina. H&E stained elephant retina. (A) Specimens containing nuclei within the OPL. Svc11-24, svc11-10, svc11-23, svc11-09, svc11-22, svc11-20, svc11-19, svc11-13, svc11-14, svc11-03 (left to right). (A) Specimens without nuclei in OPL. Svc11-11, svc11-07, svc11-15, svc11-04, svc11-06, svc11-16, svc11-02, svc11-12 (left to right). Age in years shown in top left of image. X100 magnification. Section 8μm thick.

**Figure 5:**
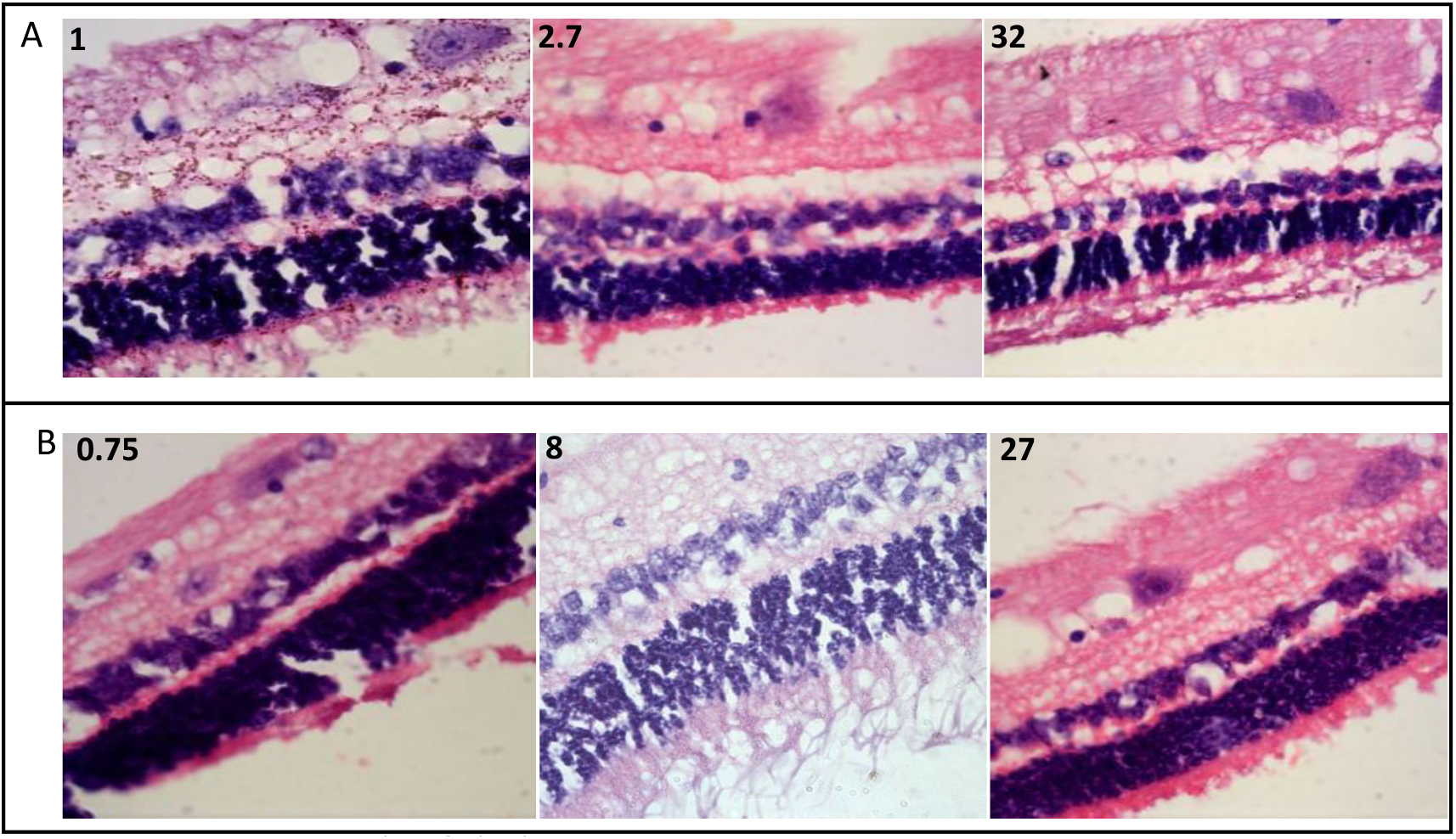
Deterioration of the inner plexiform layer was observed in a few specimens. H&E micrographs of elephant retina. (A) Specimens with deterioration of the IPL. Svc11-10, svc11-09, svc11-12 (left to right). (B) No deterioration was seen in other specimens IPL. Svc11-11, svc11-04, svc11-02 (left to right). Age in years shown in top left of image. X63 magnification. Paraffin section cut at 8μm.

**Figure 6:**
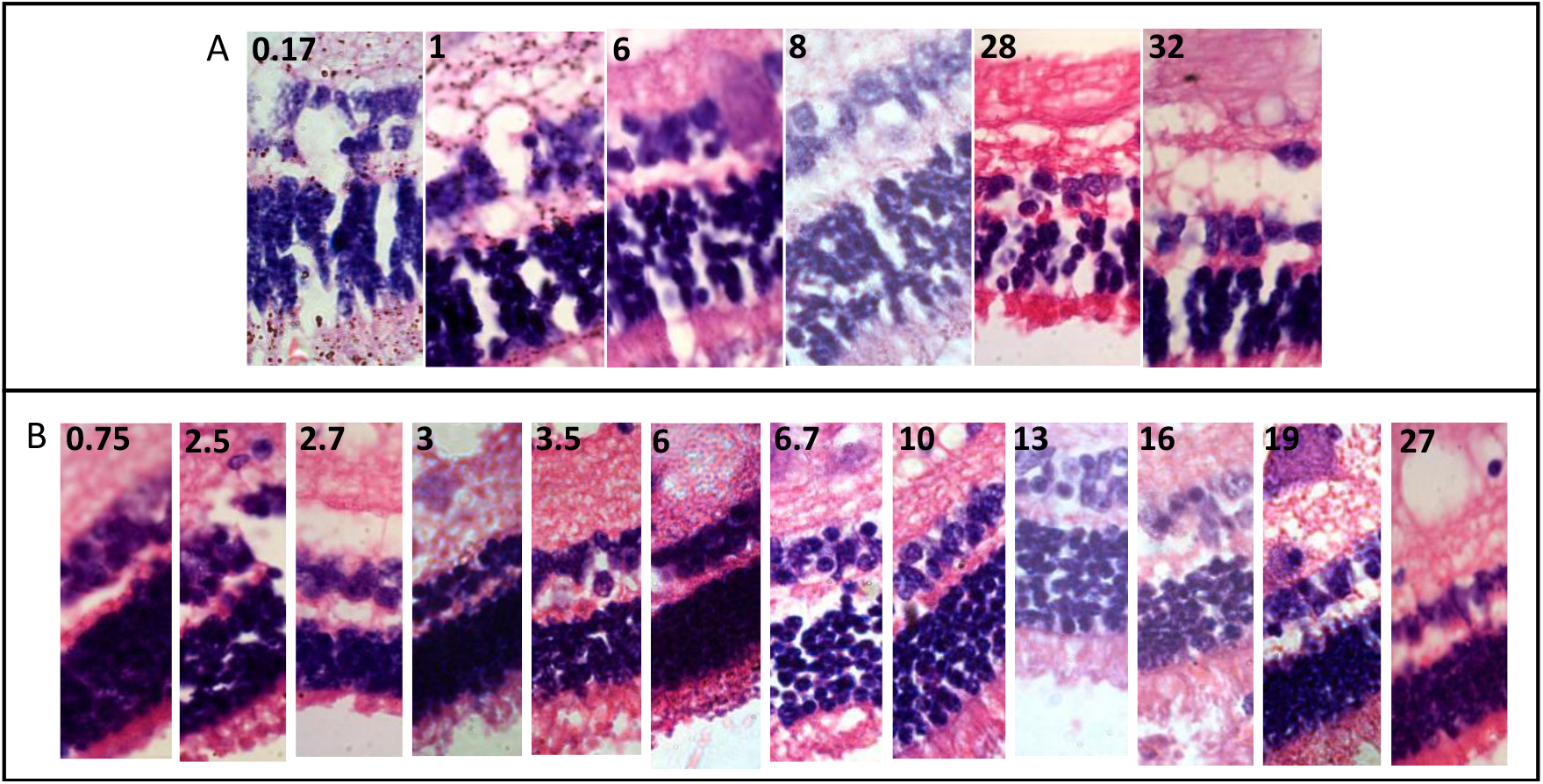
Columns of nuclei were missing in the ONL of several specimens. H&E stained sections. Sections where gaps were observed in the ONL. Svc11-24, svc11-10, svc11-07, svc11-04, svc11-03, svc11-12 (left to right). (B) For comparison, samples where nuclei and uniform in ONL. Svc11-11, svc11-23, svc11-09, svc11-22, svc11-20, svc11-15, svc11-19, svc11-06, svc11-13, svc11-14, svc11-16, svc11-02 (left to right). Age in years shown in top left of image. X100 magnification. Section is 8μm thick.

**Figure 7:**
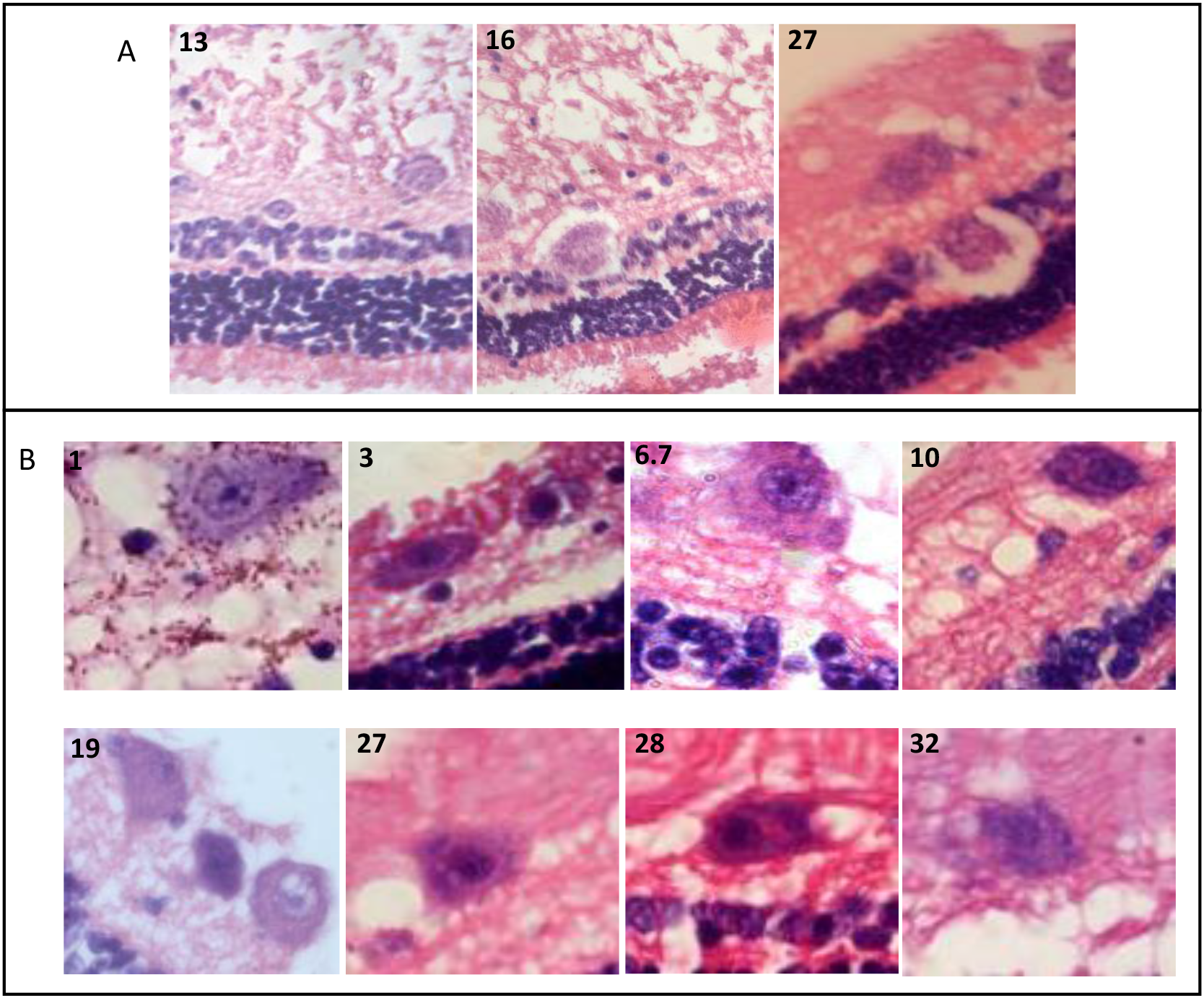
Giant ganglia and another large cell were observed in the retinas of a portion of samples. H&E stained micrographs of elephant retina. Slides showing unidentified cells. X40 magnification. Svc11-13, svc11-14, svc11-02 (left to right). Slides showing giant ganglia cells. X63 magnification. Svc11-10, svc11-22, svc11-19, svc11-06, svc11-16, svc11-02, svc11-03, svc11-12 (left to right). Age in years shown in top left of image. Paraffin section cut at 8μm.

The oldest eye had less well defined nuclei and ganglia than the younger animals (figures 4, 5 and 7). Similar changes occur in human senescent cells, which have disrupted cell and nuclear structure, and changes in chromatin affecting the nuclear pigment (Oberdoerffer and Sinclair, 2007; Shin et al., 2010). This also supports the possibility that the thickening ONL observed was related to swelling of senescent nuclei (Kobayashi et al., 2008). Elephants have been reported to have giant RGCs within their retina, which are present in in younger animals as well, demonstrated in figure 7 and Kuhrt et al. (2017). It was thought the cells within the INL (figure 7A) were also giant ganglia cells, as they can often be found in that layer (Kuhrt et al., 2017), but they were larger and less densely stained to those seen in Pettigrew et al., (2010).

The African elephant samples used in this study had the largest range of ages assessed to date in a study of eyes in this species. Our study supports previous findings on tapetal colour, eye diameter changes with age, and the presence of giant ganglia within the elephant retina. Some deviations from previous literature were also observed, relating to a change in retinal layers with age and the presence of abnormal cells in the INL, which could indicate possible age-related degeneration of the eye. This study has provided a survey of retinal structure in the African elephant across a large range of ages; in the future, a more detailed investigation of the cellular organisation of the elephant retina would be of interest to describe their species specific RGC clusters and photoreceptor mosaics. There is clearly much that remains to be learned about the eyes of African elephants.

## Acknowledgements

This study was funded by the School of Veterinary Medicine and Science at the University of Nottingham. TLI was supported by the Biotechnology and Biological Sciences Research Council [grant number BB/J014508/1],

---

**Supplemental table 1.**
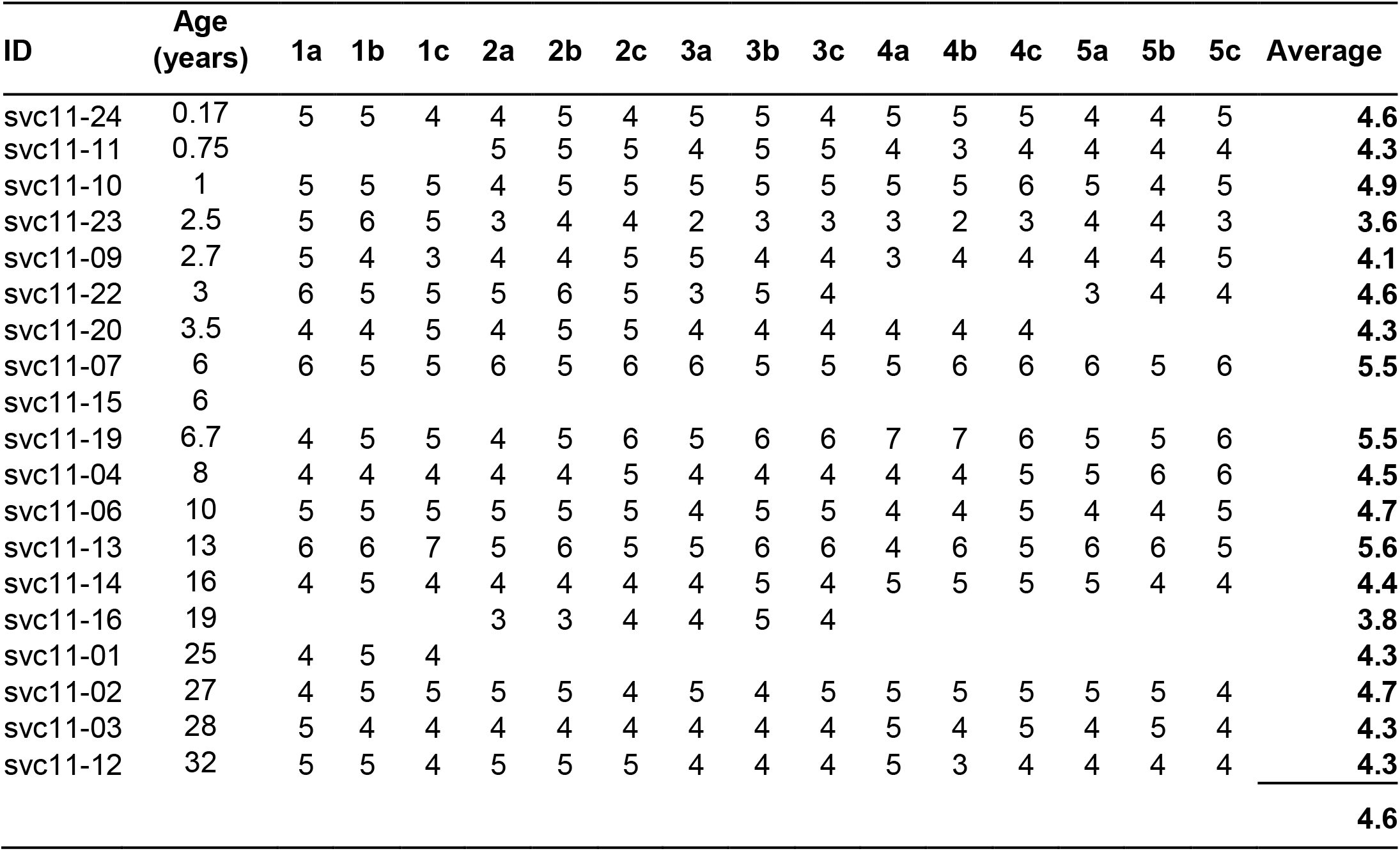
Photoreceptor count from the ONL in each of the 5 H&E slides. The overall average was 4.6. All eyes containing retina are shown, though counts were not taken from all slides due to overstaining. From an unpaired t-test, we suggest that there is no significant difference in the number of photoreceptor nuclei within a vertical column between juvenile and adult *L. africana* retinas (p= 0.17)

## Supplemental figures

**Supplemental figure 2:**
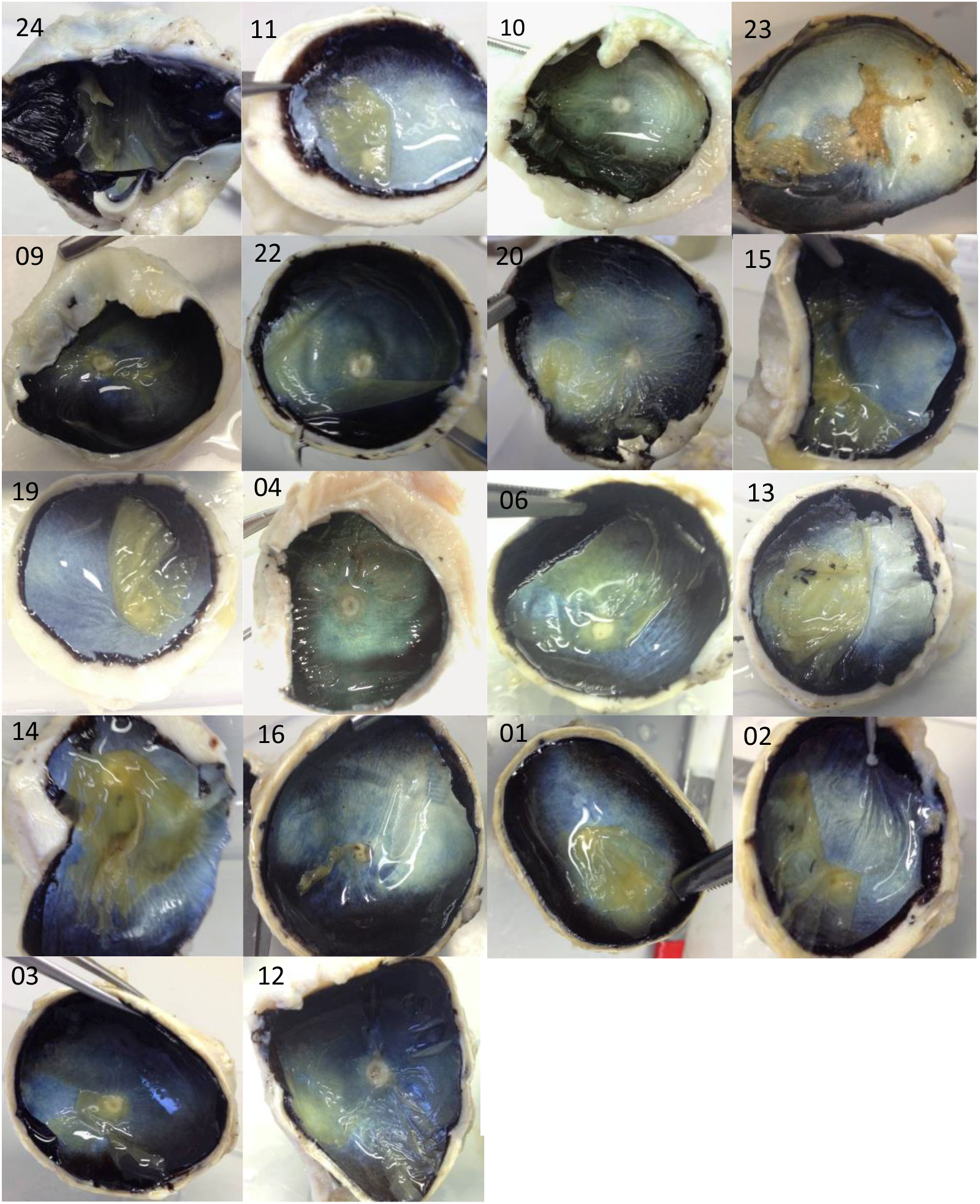
Photographs of the posterior eye from youngest to oldest (left to right). svc11-ID numbers shown. There are no image svc11-07 and −18

**Supplemental figure 2:**
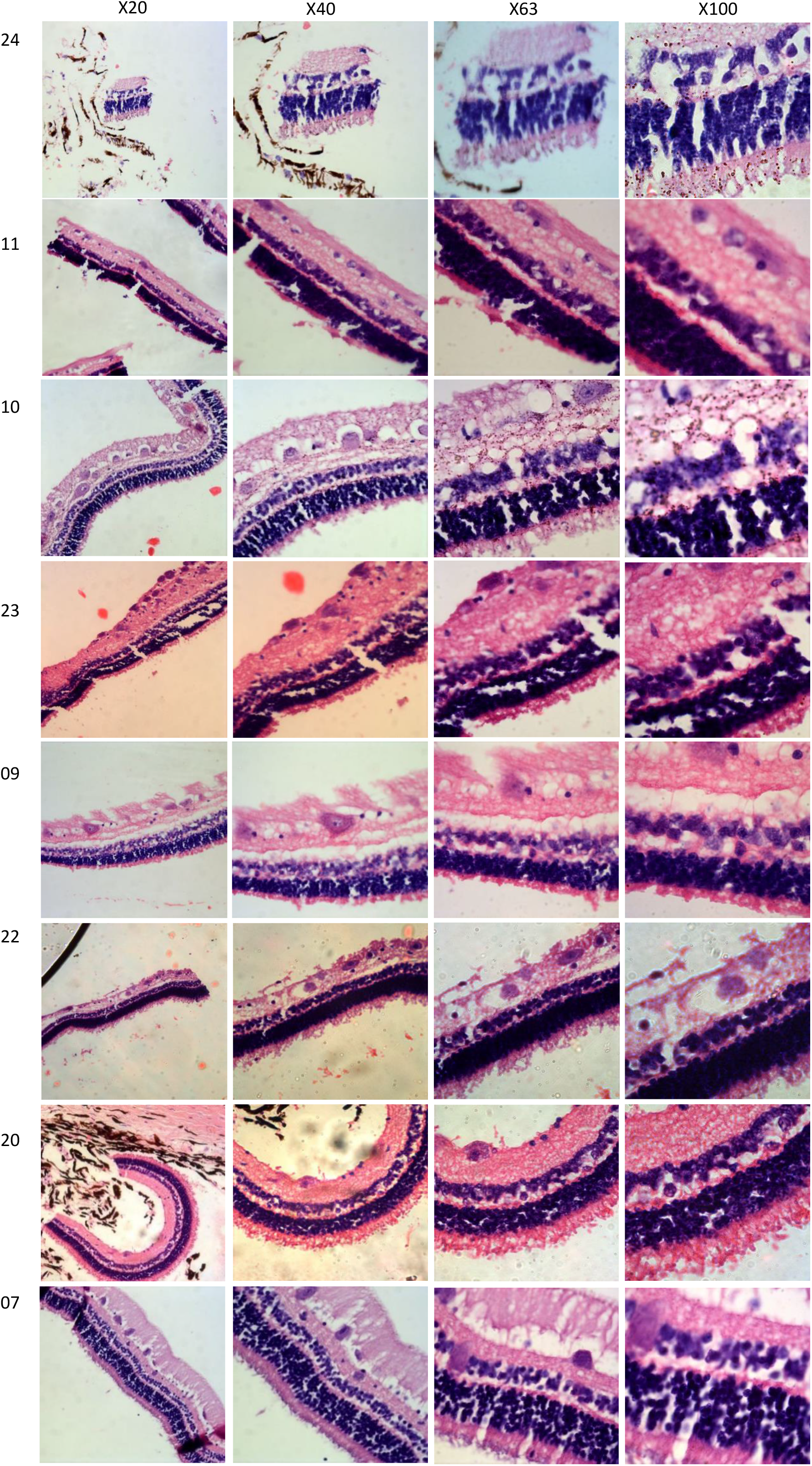
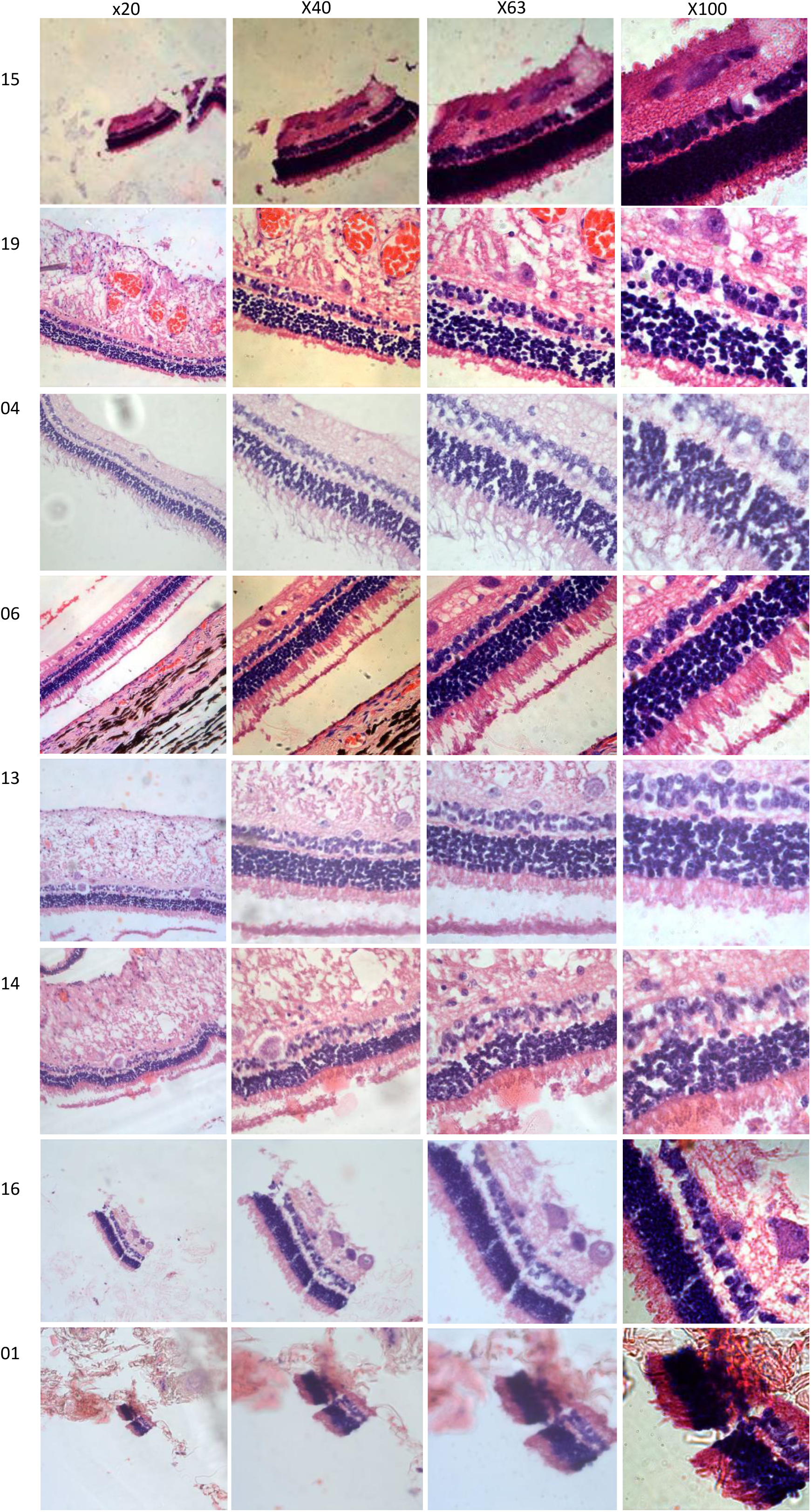
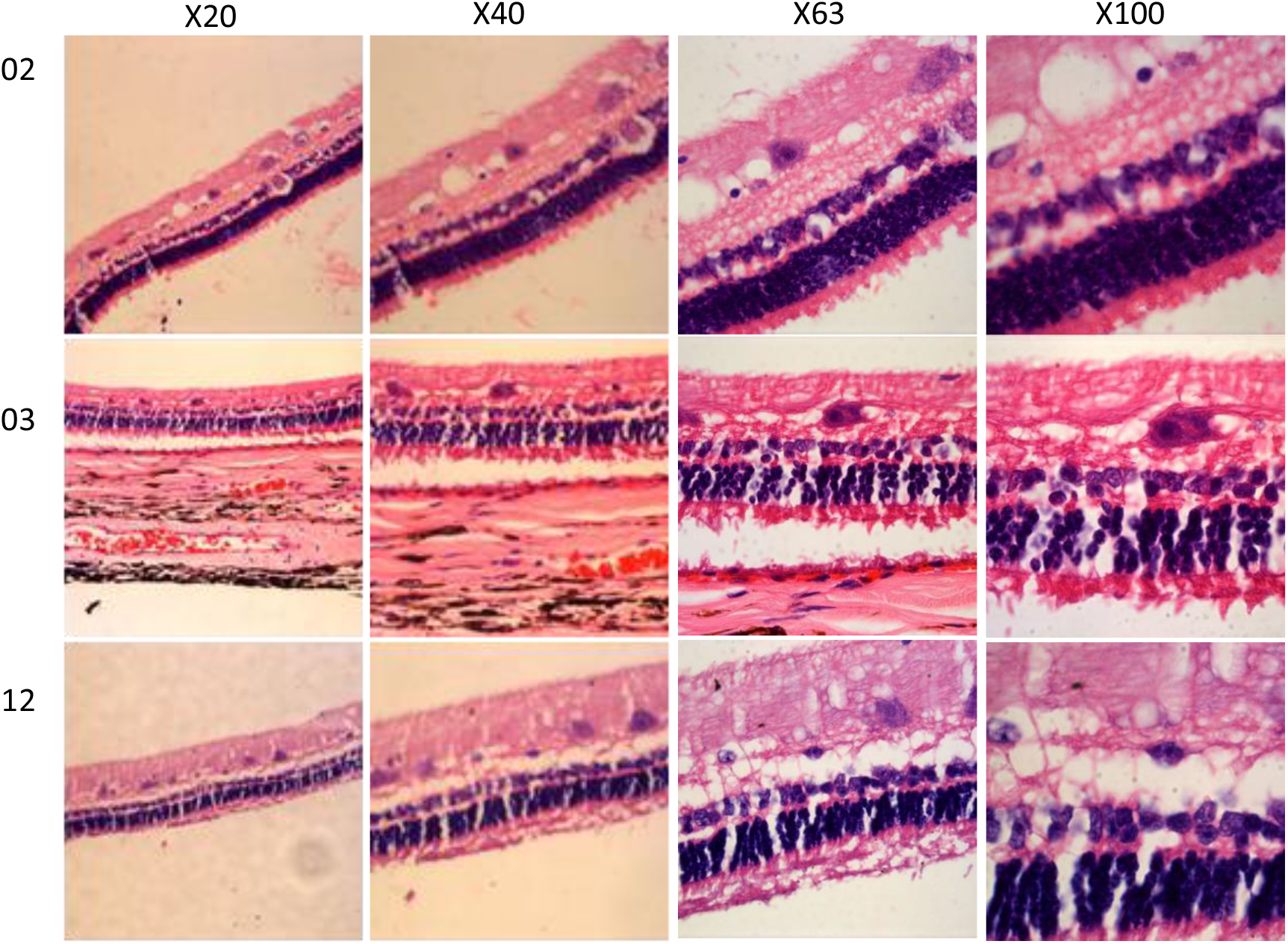
Light micrographs of the elephant eye retinas at x20, x40, x63, and x100 magnifications. Svc11-ID numbers shown. Retinas were not found on svc11-05, −12, −17, −18, −21. H&E stain. Paraffin section cut at 8μm.

## Bibliography

Bapodra, P., Bouts, T., Mahoney, P., Turner, S., Silva-Fletcher, A., and Waters, M. (2010). Ultrasonographic anatomy of the Asian elephant (Elephas maximus) eye. J. Zoo Wildl. Med. 41, 409–417. doi:10.1638/2009-0018.1.

Blagosklonny, M. V. (2006). Aging and immortality: Quasi-programmed senescence and its pharmacologic inhibition. Cell Cycle 5, 2087–2102. doi:10.4161/cc.5.18.3288.

Blagosklonny, M. V. (2012). Answering the ultimate question “What is the proximal cause of aging?” Aging (Albany. NY). 4, 861–877. doi:10.18632/aging.100525.

Bonnel, S., Mohand-Said, S., and Sahel, J. A. (2003). The aging of the retina. Exp. Gerontol. 38, 825–831. doi:10.1016/S0531-5565(03)00093-7.

Campisi, J. (2013). Aging, cellular senescence, and cancer. Annu. Rev. Physiol. 75, 685–705. doi:10.1146/annurev-physiol-030212-183653.

Curcio, C. A., Millican, C. L., Allen, K. A., and Kalina, R. E. (1993). Aging of the human photoreceptor mosaic: Evidence for selective vulnerability of rods in central retina. Investig. Ophthalmol. Vis. Sci. 34, 3278–3296.

Dawson, W. W., Hawthorne, M. N., Jenkins, R. L., and Goldston, R. T. (1982). Giant neural systems in the inner retina and optic nerve of small whales. J. Comp. Neurol. 205, 1–7. doi:10.1002/cne.902050102.

Ehrenhofer, M. C. A., Deeg, C. A., Reese, S., Liebich, H. G., Stangassinger, M., and Kaspers, B. (2002). Normal structure and age-related changes of the equine retina. Vet. Ophthalmol. 5, 39–47. doi:10.1046/j.1463-5224.2002.00210.x.

El-Sayyad, H. I. H., Khalifa, S. A., Al-Gebaly, A. S., and El-Mansy, A. A. (2014). Aging related changes of retina and optic nerve of uromastyx aegyptia and falco tinnunculus. ACS Chem. Neurosci. 5, 39–50. doi:10.1021/cn400154k.

Eliasieh, K., Liets, L. C., and Chalupa, L. M. (2007). Cellular reorganization in the human retina during normal aging. Investig. Ophthalmol. Vis. Sci. 48, 2824–2830. doi:10.1167/iovs.06-1228.

Eorge L Indsay, G., and Ohnson, J. (1901). I. Contributions to the comparative anatomy of the mammalian eye, chiefly based on ophthalmoscopic examination. Philos. Trans. R. Soc. London. Ser. B, Contain. Pap. a Biol. Character 194, 1–82. doi:10.1098/rstb.1901.0001.

Fischer, A. H., Jacobson, K. A., Rose, J., and Zeller, R. (2008). Hematoxylin and eosin staining of tissueand cell sections. Cold Spring Harb. Protoc. 3, pdb.prot4986. doi:10.1101/pdb.prot4986.

Friedlander, M. (2007). Fibrosis and diseases of the eye. J. Clin. Invest. 117, 576–586. doi:10.1172/JCI31030.

Gao, H., and Hollyfield, J. G. (1992). Aging of the human retina: Differential loss of neurons and retinal pigment epithelial cells. Investig. Ophthalmol. Vis. Sci. 33, 1–17.

Gonzalez-Freire, M., De Cabo, R., Bernier, M., Sollott, S. J., Fabbri, E., Navas, P., et al. (2015). Reconsidering the Role of Mitochondria in Aging. Journals Gerontol. -Ser. A Biol. Sci. Med. Sci. doi:10.1093/gerona/glv070.

Howland, H. C., Merola, S., and Basarab, J. R. (2004). The allometry and scaling of the size of vertebrate eyes. Vision Res. 44, 2043–2065. doi:10.1016/j.visres.2004.03.023.

Kobayashi, Y., Sakemura, R., Kumagai, A., Sumikawa, E., Fujii, M., and Ayusawa, D. (2008). Nuclear swelling occurs during premature senescence mediated by MAP kinases in normal human fibroblasts. Biosci. Biotechnol. Biochem. 72, 1122–1125. doi:10.1271/bbb.70760.

Kuhrt, H., Bringmann, A., Härtig, W., Wibbelt, G., Peichl, L., and Reichenbach, A. (2017). The retina of asian and African elephants: Comparison of newborn and adult. Brain. Behav. Evol. 89, 84–103. doi:10.1159/000464097.

Kuhrt, H., Gryga, M., Wolburg, H., Joffe, B., Grosche, J., Reichenbach, A., et al. (2012). Postnatal mammalian retinal development: Quantitative data and general rules. Prog. Retin. Eye Res. 31, 605–621. doi:10.1016/j.preteyeres.2012.08.001.

Kyosseva, S. V. (2016). Targeting MAPK Signaling in Age-Related Macular Degeneration. Ophthalmol. Eye Dis. 8, OED.S32200. doi:10.4137/oed.s32200.

L?pez-Ot?n, C., Blasco, M. A., Partridge, L., Serrano, M., and Kroemer, G. (2013). The Hallmarks of Aging. Cell 153, 1194–1217. doi:10.1016/j.cell.2013.05.039.

Lee, M. W., Kim, J. mi, Lim, H. Bin, Shin, Y. Il, Lee, Y. H., and Kim, J. Y. (2020). Longitudinal Changes in Ganglion Cell–Inner Plexiform Layer of Fellow Eyes in Unilateral Neovascular Age-Related Macular Degeneration. Am. J. Ophthalmol. 212, 17–25. doi:10.1016/j.ajo.2019.12.003.

Lee, P. C., Sayialel, S., Lindsay, W. K., and Moss, C. J. (2012). African elephant age determination from teeth: Validation from known individuals. Afr. J. Ecol. 50, 9–20. doi:10.1111/j.1365-2028.2011.01286.x.

Li, M., Huisingh, C., Messinger, J., Dolz-Marco, R., Ferrara, D., Bailey Freund, K., et al. (2018). Histology of geographic atrophy secondary to age-related macular degeneration a multilayer approach. Retina 38, 1937–1953. doi:10.1097/IAE.0000000000002182.

Liets, L. C., Eliasieh, K., Van Der List, D. A., and Chalupa, L. M. (2006). Dendrites of rod bipolar cells sprout in normal aging retina. Proc. Natl. Acad. Sci. U. S. A. 103, 12156–12160. doi:10.1073/pnas.0605211103.

Life expectancy at birth (2015). doi:10.1787/how_life-2015-graph25-en.

Milholland, B., Suh, Y., and Vijg, J. (2017). Mutation and catastrophe in the aging genome. Exp. Gerontol. 94, 34–40. doi:10.1016/j.exger.2017.02.073.

Müller, B., Butz, E., Peichl, L., and Haverkamp, S. (2013). The rod pathway of the microbat retina has bistratified rod bipolar cells and tristratified AII amacrine cells. J. Neurosci. 33, 1014–1023. doi:10.1523/JNEUROSCI.2072-12.2013.

Mullins, R. F., and Skeie, J. M. (2010). Essentials of retinal morphology. Neuromethods 46, 1–11. doi:10.1007/978-1-60761-541-5_1.

Murphy, C. J., Kern, T. J., and Howland, H. C. (1992). Refractive state, corneal curvature, accommodative range and ocular anatomy of the Asian elephant (Elephas maximus). Vision Res. 32, 2013–2021. doi:10.1016/0042-6989(92)90062-N.

Newsome, D. A., Huh, W., and Green, W. R. (1987). Bruch’s membrane age-related changes vary by region. Curr. Eye Res. 6, 1211–1221. doi:10.3109/02713688709025231.

Nussenblatt, R. B., Kaufman, S. C., Palestine, A. G., Davis, M. D., and Ferris, F. L. (1987). Macular Thickening and Visual Acuity: Measurement in Patients with Cystoid Macular Edema. Ophthalmology 94, 1134–1139. doi:10.1016/S0161-6420(87)33314-7.

Oberdoerffer, P., and Sinclair, D. A. (2007). The role of nuclear architecture in genomic instability and ageing. Nat. Rev. Mol. Cell Biol. 8, 692–702. doi:10.1038/nrm2238.

Ollivier, F. J., Samuelson, D. A., Brooks, D. E., Lewis, P. A., Kallberg, M. E., and Komáromy, A. M. (2004). Comparative morphology of the tapetum lucidum (among selected species). Vet. Ophthalmol. 7, 11–22. doi:10.1111/j.1463-5224.2004.00318.x.

Pettigrew, J. D., Bhagwandin, A., Haagensen, M., and Manger, P. R. (2010). Visual acuity and heterogeneities of retinal ganglion cell densities and the tapetum lucidum of the African elephant (Loxodonta africana). Brain. Behav. Evol. 75, 251–261. doi:10.1159/000314898.

Ramrattan, R. S., Van der Schaft, T. L., Mooy, C. M., De Bruijn, W. C., Mulder, P. G. H., and De Jong, P. T. V. M. (1994). Morphometric analysis of Bruch’s membrane, the choriocapillaris, and the choroid in aging. Investig. Ophthalmol. Vis. Sci. 35, 2857–2864.

Samuel, M. A., Zhang, Y., Meister, M., and Sanes, J. R. (2011). Age-related alterations in neurons of the mouse retina. J. Neurosci. 31, 16033–16044. doi:10.1523/JNEUROSCI.3580-11.2011.

Shin, D. M., Kucia, M., and Ratajczak, M. Z. (2010). Nuclear and chromatin reorganization during cell senescence and aging – A mini-review. Gerontology 57, 76–84. doi:10.1159/000281882.

Stoeger, A. S., Zeppelzauer, M., and Baotic, A. (2014). Age-group estimation in free-ranging African elephants based on acoustic cues of low-frequency rumbles. Bioacoustics 23, 231–246. doi:10.1080/09524622.2014.888375.

Stone, J., and Halasz, P. (1989). Topography of the retina in the elephant loxodonta Africana. Brain. Behav. Evol. 34, 84–95. doi:10.1159/000116494.

Sugita, Y., Yamamoto, H., Maeda, Y., and Furukawa, T. (2020). Influence of Aging on the Retina and Visual Motion Processing for Optokinetic Responses in Mice. Front. Neurosci. 14, 1255. doi:10.3389/fnins.2020.586013.

Szilard, L. (1959). ON THE NATURE OF THE AGING PROCESS. Proc. Natl. Acad. Sci. 45, 30–45. doi:10.1073/pnas.45.1.30.

Vecino, E., Hernández, M., and García, M. (2004). Cell death in the developing vertebrate retina. Int. J. Dev. Biol. 48, 965–974. doi:10.1387/ijdb.041891ev.

Wang, S., Zheng, Y., Li, Q., He, X., Ren, R., Zhang, W., et al. (2020). Deciphering primate retinal aging at single-cell resolution. Protein Cell, 1–10. doi:10.1007/s13238-020-00791-x.

Werner, M., Chott, A., Fabiano, A., and Battifora, H. (2000). Effect of formalin tissue fixation and processing on immunohistochemistry. Am. J. Surg. Pathol. 24, 1016–1019. doi:10.1097/00000478-200007000-00014.

